# Towards a human brain EV atlas: Characteristics of EVs from different brain regions, including small RNA and protein profiles

**DOI:** 10.1101/2023.05.06.539665

**Authors:** Yiyao Huang, Tanina Arab, Ashley E. Russell, Emily R. Mallick, Rajini Nagaraj, Evan Gizzie, Javier Redding-Ochoa, Juan C. Troncoso, Olga Pletnikova, Andrey Turchinovich, David A. Routenberg, Kenneth W. Witwer

**Affiliations:** Department of Molecular and Comparative Pathobiology, Johns Hopkins University School of Medicine, Baltimore, MD, USA; Department of Biology, School of Science, Penn State Erie, The Behrend College, Erie, PA, United States; Meso Scale Diagnostics, LLC, Rockville, MD, USA; Department of Pathology, Johns Hopkins University School of Medicine, Baltimore, MD, USA; Department of Neurology, Johns Hopkins University School of Medicine, Baltimore, MD, USA; Department of Pathology and Anatomical Sciences, Jacobs School of Medicine and Biomedical Sciences, University at Buffalo, Buffalo, NY, USA; Division of Cancer Genome Research, German Cancer Research Center DKFZ, Heidelberg, Germany; Heidelberg Biolabs GmbH, Mannheim, Germany; The Richman Family Precision Medicine Center of Excellence in Alzheimer’s Disease, Johns Hopkins University School of Medicine, Baltimore, MD, US

**Keywords:** Brain, Extracellular vesicles, ectosomes, exosomes, tissue, corpus callosum, thalamus, orbitofrontal, postcentral gyrus, occipital gyrus, cerebellum, medulla, hippocampus

## Abstract

Extracellular vesicles (EVs) are released from different cell types in the central nervous system (CNS) and play roles in regulating physiological and pathological functions. Although brain-derived EVs (bdEVs) have been successfully collected from brain tissue, there is not yet a “bdEV atlas” of EVs from different brain regions. To address this gap, we separated EVs from eight anatomical brain regions of a single individual and subsequently characterized them by count, size, morphology, and protein and RNA content. The greatest particle yield was from cerebellum, while the fewest particles were recovered from the orbitofrontal, postcentral gyrus, and thalamus regions. EV surface phenotyping indicated that CD81 and CD9 were more abundant than CD63 for all regions. Cell-enriched surface markers varied between brain regions. For example, putative neuronal markers NCAM, CD271, and NRCAM were more abundant in medulla, cerebellum, and occipital regions, respectively. These findings, while restricted to tissues from a single individual, suggest that additional studies are merited to lend more insight into the links between EV heterogeneity and function in the CNS.

## Introduction

Extracellular vesicles (EVs) are a diversity of membranous, cell-released particles that are involved in a wide range of processes by shuttling biological materials out of and between cells^1^. In the central nervous system (CNS), EVs are released from different CNS cell types^2^, including neurons and glia. These CNS EVs regulate physiological functions in the CNS like neuronal firing, synaptic plasticity, and myelin sheath maintenance^3^, and also exert pathological functions in neurodegenerative diseases by spreading neuroinflammatory factors and toxic protein aggregates^4–6^. EVs found in brain tissue, e.g., in interstitial fluid or associated with cells or extracellular matrix, are termed brain-derived EVs (bdEVs)^5,7–10^. bdEVs are increasingly studied since they may lend insights into CNS disease mechanisms and may also betray disease when released into easily accessed biological fluids ^11–15^.

The brain can be divided into anatomical regions with diversity of function and cell composition. Neurons, astrocytes, microglia, oligodendrocytes, and other cells vary across regions in number, density, morphology, and molecular signature^16–18^. For example, pyramidal neurons are prominent in the cerebral cortex, while granule and Purkinje cells are found only in the cerebellum^19–21^, and microglia are more abundant and dense in cortical regions compared with cerebellum^17^. Microglia from different brain regions are also affected differently in aging^22,23^. Regional differences in disease progression are also reported. For example, Alzheimer ‘s disease (AD) affects the hippocampus (HP) and entorhinal cortex (ENT) more severely and earlier than other regions^24^. To capture information that might be missed by single-region analysis, several studies have profiled multiple brain tissue regions, including the Allen Brain Atlas data portal^25^ (www.brain-map.org), a study of two brain regions in schizophrenia^26^ (http://eqtl.brainseq.org/phase2), and protein profiling of six brain regions in AD and asymptomatic controls^24^ (www.manchester.ac.uk/dementia-proteomes-project).

In this study, we asked the question of whether bdEVs might also display brain region-specific signatures. We separated bdEVs from eight brain regions: orbitofrontal gyrus (ORB), postcentral gyrus (POSTC), hippocampus (HIPPO), thalamus (THAL), occipital gyrus (OCC), medulla (MED), corpus callosum (CORP), and cerebellum (CBLM). bdEVs were thoroughly characterized to gather any evidence of regional differences.

## MATERIALS AND METHODS

### Tissue collection and preparation

Human postmortem brain tissue samples were obtained from the Brain Resource Center (Department of Pathology, Johns Hopkins University School of Medicine) following brain autopsy with complete neuropathological examination. The brain of a human male was dissected by neuroanatomical region. Sections were stored at −80°C prior to bdEV separation.

### Brain extracellular vesicle separation

To avoid batch effects and operator bias, EVs were separated from all tissue regions simultaneously. Separation was as previously described^10^. Approximately 20 mg of each tissue region was dissected and stored at −80°C for Western blot (WB) (brain homogenate, BH). The remaining frozen tissue was weighed and gently digested using collagenase type 3 enzyme in Hibernate-E solution for 20 min at 37°C. A solution containing 1X PhosSTOP (Sigma, 4906837001) and Complete Protease Inhibitor solution (Sigma,11697498001) (PI/PS) was added to stop the enzymatic reaction. Differential centrifugation was performed at 4°C. Dissociated tissue was centrifuged at 300 × g for 10 min. The supernatant was pipetted off using a 10 ml serological pipette, transferred to a fresh tube, and centrifuged at 2000 × g for 15 min. Pellets were collected as a “2K” fraction. The supernatant was further depleted of debris and large bodies through a gentle 0.22-μm filtration at a slow flow rate of 5 ml/min. The filtered material was centrifuged at 10,000 x g for 30 min using swinging-bucket rotor TH-641 (Thermo Scientific, k-factor 114, acceleration and deceleration settings of 9). Pellets (“10K “) were resuspended in 150 μl PBS containing 1X PI/PS by pipetting up and down ten times, vortexing for 30 seconds, and incubating on ice for 20 minutes, followed by a repeat of the pipetting and vortexing steps. Resuspended 10K bdEVs were aliquoted and stored at -80°C. Supernatants were transferred into 100 kilodalton (kDa) molecular weight cut-off (MWCO) protein concentrators (Millipore-Sigma, UFC 805024) and concentrated from 5 ml to 0.5 ml. Retentate was then applied to the top of qEV Original size exclusion chromatography (SEC) columns (Izon Science, SP1, 70 nm) that were pre-rinsed with 15 ml PBS. 0.5-ml fractions were collected by elution with PBS, using Izon automated fraction collectors (AFCs; Izon Science, Cambridge, MA). Fractions 1-6 (3 ml total) were considered the void volume; fractions 7-10 were pooled as EV-enriched fractions. EV-enriched fractions were transferred to polypropylene UC tubes and centrifuged at 100,000 × g for 70 min, using the TH-641 swinging-bucket rotor as described above. The supernatant was poured off, and UC tubes were placed upright and inverted on a piece of tissue to drain the residual buffer. Pellets (“100K,/ bdEVs) were resuspended in 120 μl PBS (1X PI/PS) using the same protocol as for “10K/. Aliquots were stored at -80°C.

### Nano-flow cytometry measurement (NFCM)

Particle concentration and size profiles of bdEVs were assessed by nano-flow (Flow NanoAnalyzer, NanoFCM, Nottingham, England) as described previously ^10,27^. Instrument calibrations for concentration and size distribution were done using reference 200-nm polystyrene beads and a silica nanosphere mixture (diameters of 68, 91, 113, and 151 nm), respectively. 2 μl of bdEV resuspension was used for serial dilutions from 1:100 to 1:200 in DPBS to determine the best particle count range as required by the manual and as reported previously ^27^, and events were recorded for 1 min. The particle concentration and size of particles were converted by calibration curves based on flow rate and side-scatter intensity. Washing steps were performed using a cleaning solution to avoid contamination across samples.

### Transmission electron microscopy (TEM)

bdEVs were imaged by TEM as previously described ^27^. Briefly, 10 μl of each sample was freshly thawed and adsorbed to glow-discharged carbon-coated 400 mesh copper grids by flotation for 2 min. Three consecutive drops of 1× Tris-buffered saline were prepared on Parafilm. Grids were washed by moving from one drop to another, with a flotation time of 10 seconds on each drop. The rinsed grids were then negatively stained with 1% uranyl acetate (UAT) with tylose (1% UAT in deionized water (dIH2O), double-filtered through a 0.22-μm filter). Grids were blotted, then excess UAT was aspirated, leaving a thin layer of stain. Grids were imaged on a Hitachi 7600 TEM operating at 80 kV with an XR80 charge-coupled device (8 megapixels, AMT Imaging, Woburn, MA, USA).

### Western blot (WB)

BH with collagenase (BH), 2K, 10K, bdEVs and SEC protein fractions were lysed in 1× radioimmunoprecipitation assay buffer (RIPA, cell signaling #9806) supplemented with a protease inhibitor cocktail. Samples were loaded as equal volumes of 20 μl and were resolved using a 4% to 15% Criterion TGX Stain-Free Precast gel, then transferred onto an Immuno-Blot PVDF membrane. Antibodies to CD63, and CD9 (BD Pharmingen #556019 and BioLegend #312102) were used to detect EV membrane markers, anti-alix (ab186429) for detection of an EV internal protein, anti-calreticulin antibody (cell signaling #12238)) was used to detect endoplasmic reticulum contamination. Primary antibodies were diluted 1:1000 in PBS-T containing 5% blotting-grade blocker (Bio-Rad, #1706404). Membranes were incubated overnight (≈16 h). After several washes in PBS-T, rabbit anti-mouse IgGk BP-HRP and mouse anti-rabbit IgGk BP-HRP secondary antibodies (Santa Cruz #516102 and #sc-2357 respectively) were diluted 1:5000 in blocking buffer, and membranes were incubated for 1 h at room temperature (RT). SuperSignal West Pico PLUS Chemiluminescent Substrate (Thermo Fisher, 34580) was applied, and blots were visualized using a Thermo Fisher iBright 1500 imaging system.

### Single-particle interferometric reflectance imaging sensor (SP-IRIS)

EVs were phenotyped with EV-TETRA-C ExoView Tetraspanin kits and an ExoView TMR100 scanner (NanoView Biosciences, Boston, MA) according to the manufacturer ‘s instructions and as described previously ^27^. A total of 10 μL bdEVs were diluted in 35 μL incubation buffer (IB). 45μl of the mixture was placed and incubated with the at RT for 16h. Chips were then washed with IB and incubated with a fluorescently-labeled antibody cocktail of anti-human CD81 (JS-81, CF555), CD63 (H5C6, CF647), and CD9 (HI9a, CF488A) at dilutions of 1:1200 (v:v) in blocking solution for 1h at RT. All chips were washed and scanned with the ExoView scanner using both SP-IRIS Single Particle Interferometric Reflectance Imaging Sensor and fluorescence detection. Data were analyzed using NanoViewer 2.8.10 Software.

### Multiplexed ELISA

Prototype ultrasensitive electrochemiluminescence assays (Meso Scale Diagnostics, Rockville, MD) were used for intact EV surface marker detection. Three multiplexed assay panels were used in this study (as listed in Supplementary table 1). 5 μL of each bdEV sample was diluted 1 to 40 in MSD diluent 52 and samples were added to assay plates with capture antibody arrays and shaken continuously at RT during the EV capture step. Panel 1, comprising antibodies targeting relatively abundant surface markers, was incubated for 1 hour, while the remaining panels, targeting lower-abundance markers, were incubated for 4 hours to improve sensitivity. EVs captured by each antibody were detected using prototype MSD S-PLEX® detection reagents with a cocktail of detection antibodies targeting CD63, CD81, and CD9. Assay plates were read with MSD GOLD™ Read buffer B on an MSD® SECTOR 600 imager. ECL signal from a DPBS blank on each assay spot and ECL signal from each bdEV sample on an isotype-control capture spot were subtracted consecutively from the signal of each corresponding assay to account for non-specific binding of detector antibodies and the EVs in the sample, respectively.

### RNA extraction and small RNA sequencing

bdEV RNA was extracted by miRNeasy Mini Kit reagents (Qiagen 217004) and Zymo-Spin I Columns (Zymo Research C1003-50) according to the manufacturer ‘s instructions. bdEV RNA was resuspended in 40 μL Rnase-free water, and 8 μl was used for small RNA libraries construction by the D-Plex Small RNA-seq Kit (Diagenode C05030001). Indexes were attached using the D-Plex Single Indexes for Illumina - Set A (Diagenode C05030010) according to the manufacturer ‘s protocol. The yield and size distribution of the small RNA libraries were assessed using the Fragment Bioanalyzer™ system with DNA 1000 chip (Agilent 5067-1505). After size selection of the libraries by agarose gel cassettes (Sage Science HTG3010) on BluePippin (Sage Science) from 170-230 bp, multiplexed libraries were equally pooled to 1 nM. and prepared for deep sequencing using the NovaSeq 6000 system (Illumina) and sequenced by NovaSeq 6000 SP Reagent Kit v1.5 (100 cycles) (Illumina 20028401).

### RNA sequencing data analysis

The RNA sequencing data were analyzed as previously published ^28,29^. Original BAM files were converted into FASTQ format using Picard tools (SamToFastq command). Reads shorter than 15 nt were removed from the raw FASTQ data using cutadapt software v1.18. The reads were aligned to the custom curated human reference transcriptomes in a sequential manner using bowtie allowing 1 mismatch tolerance: trimmed and size-selected reads were mapped to RNA species with low sequence complexity and/or high number of repeats: rRNA, tRNA, RN7S, snRNA, snoRNA/scaRNA, vault RNA, RNY as well as mitochondrial chromosome (mtRNA). All reads that did not map to the above RNAs were aligned sequentially to mature miRNA, pre-miRNA, protein-coding mRNA transcripts (mRNA), and long non-coding RNAs (lncRNAs). The numbers of reads mapped to each RNA type were extracted using eXpress software based on a previous publication^30^. The mapping data was normalized using R/Bioconductor packages DESeq2^29^ and then visualized with the principal component analysis (PCA) plot. Hierarchical clustering of miRNAs was performed with Heatmapper.

### Data and methods availability

We have submitted all relevant data of our experiments to the EV-TRACK knowledgebase (EV-TRACK ID: EV230055.) ^31^. Nucleic acid sequencing data were deposited with the Gene Expression Omnibus, accession GSE226490.

## Results

### bdEV recovery and morphology characteristics were related to source brain regions

bdEV particle counts per 100 mg of tissue input were obtained by NFCM. Differences were observed between brain regions (Fig. 2a). The largest yield was from the cerebellum (CBLM), with 8.95x10e8 particles per 100 mg of tissue input, while the smallest number of particles were recovered from the orbitofrontal (ORB), postcentral gyrus (POSTC), and thalamus (THAL) regions, with concentrations ranging from 2.17x10e8 particles to 2.45x10e8 particles per 100 mg tissue. Intermediate counts were recovered from the corpus callosum (CORP), hippocampus (HIPPO), occipital gyrus (OCC), and medulla (MED). Transmission electron microscopy (TEM) showed that round to oval particles with characteristic EV morphology and size were recovered from all regions. However, differences in size distribution and morphology were observed (Fig. 2b). Overall, smaller vesicles were revealed in the bdEVs from the ORB, HIPPO, THAL, CORP, and CBLM regions, while larger vesicles were observed in the POSTC, OCC, and MED regions. Particles recovered from CBLM appeared to include a subpopulation of small particles with dense inner membrane contents. Quantification of particle size distribution by both NFCM (Fig. 2c) and TEM (Fig. 2d) confirmed that the preparations from ORB and CBLM contained the largest percentage of smaller particles, while the MED-derived population included larger particles.

**Figure 1.**
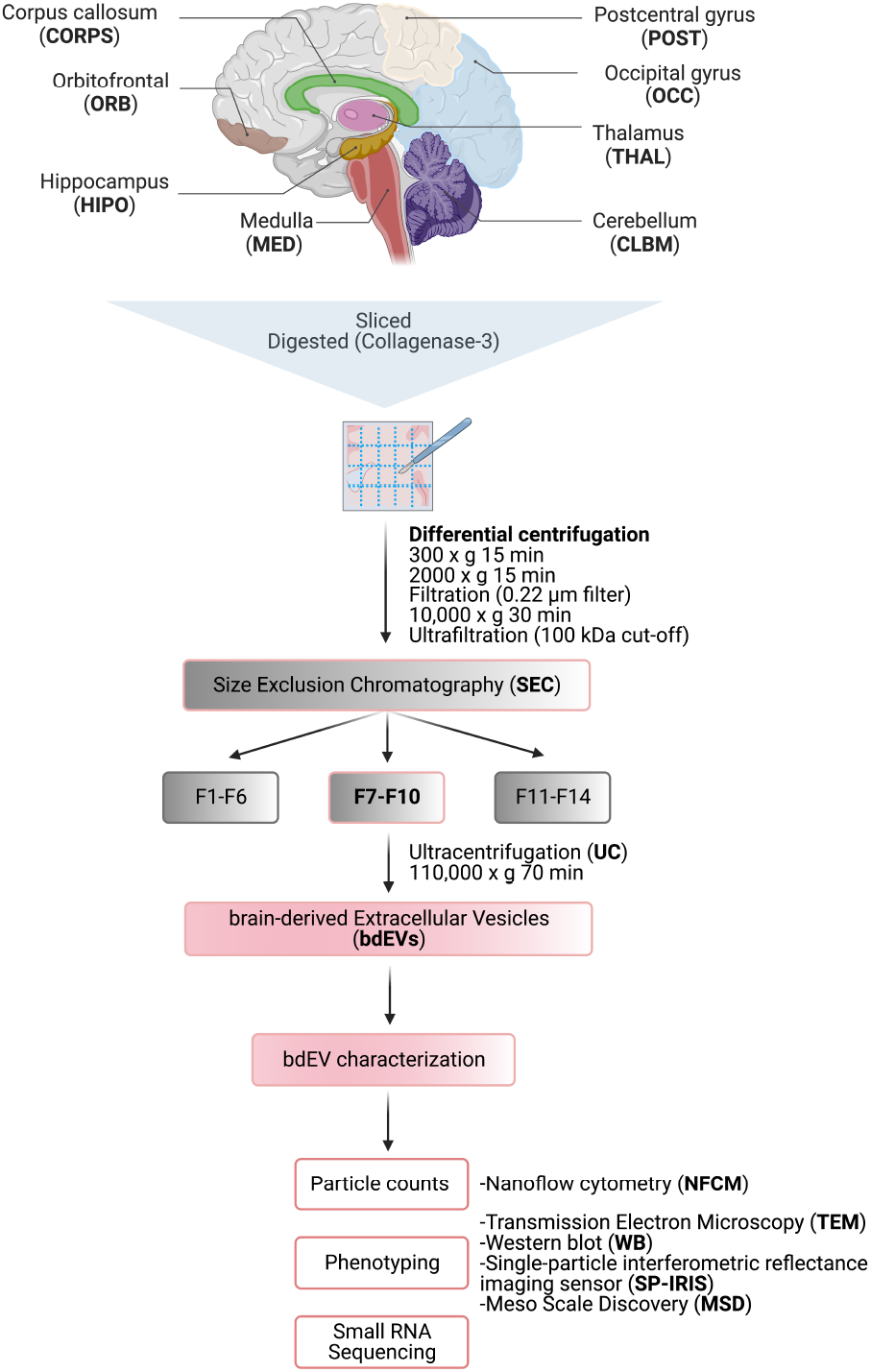
Workflow for brain-derived EV (bdEV) enrichment and characterization from different brain regions. bdEVs from 8 brain regions were separated by collagenase digestion, differential centrifugation, and size exclusion chromatography (SEC). After separation, bdEVs were characterized by particle count, imaging, protein phenotyping and small RNA sequencing. Created with BioRender.com.

**Figure 2.**
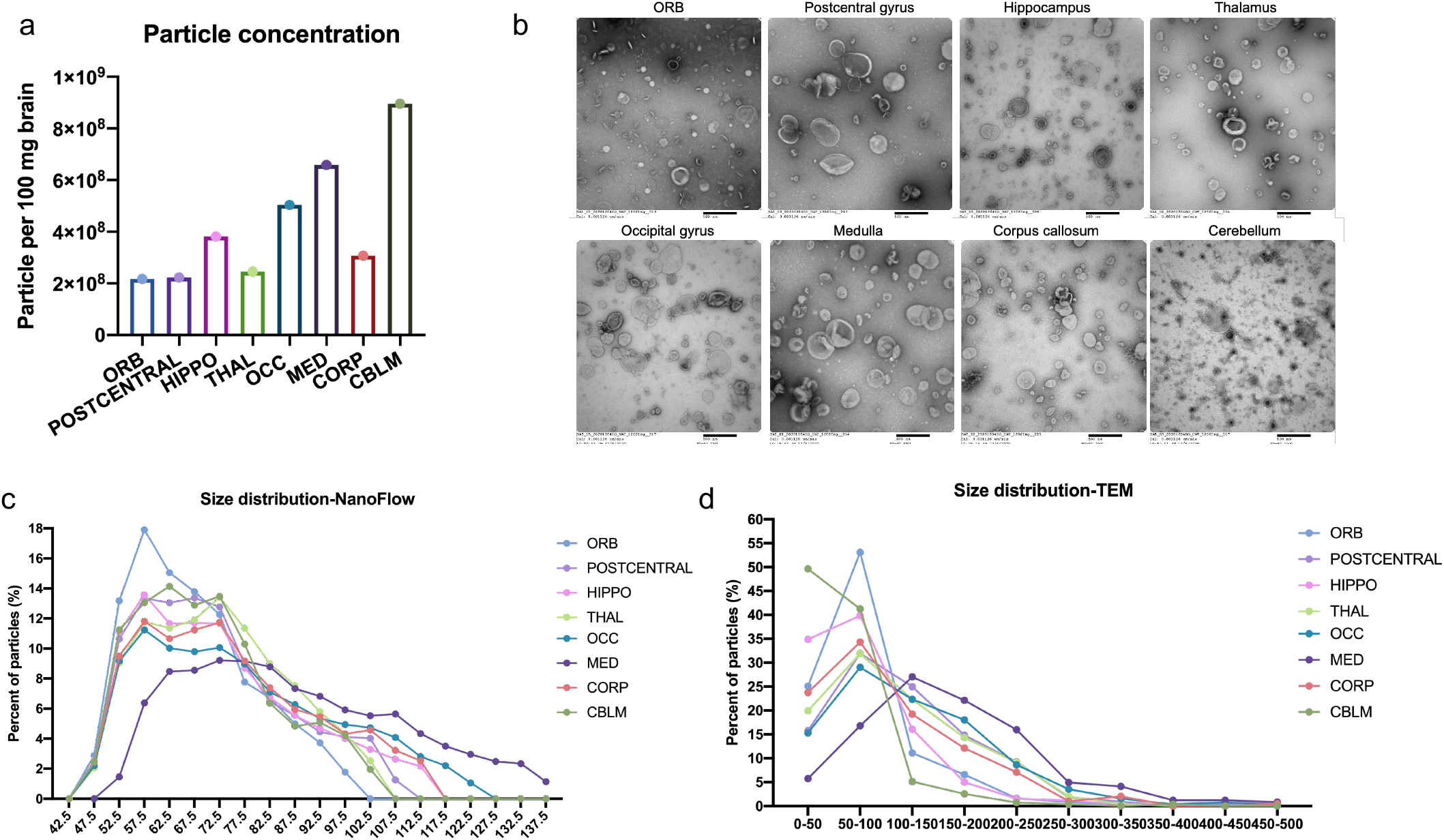
(a) Particle concentrations of bdEVs from brain regions were measured by NFCM. Particle concentration for each region was normalized by tissue mass (per 100 mg). (b) bdEVs were visualized by negative staining transmission electron microscopy (TEM) (scale bar□=□500 nm). TEM is representative of ten images taken of each region. (c) Size (diameter) distributions of bdEVs from brain regions as measured by NFCM and calculated as particles in each 5 nm size bin versus total detected particles in each sample (percentage). (d) Size distributions of bdEVs from brain regions as measured in TEM images and calculated as particles in each 50 nm size bin versus total detected particles in each sample (percentage).

### bdEV tetraspanin phenotyping

Presence of EV-enriched membrane (CD63, CD81) and cytosolic (Alix) markers, and expected EV-depleted cellular markers (calreticulin) were examined by Western blot for bdEVs from additional samples for the purpose of protocol reproducibility assessment, as well as brain homogenate (BH), 2K, and 10K (Figure S1). Abundant EV markers and depletion of cellular markers in bdEVs showed the protocol we used lead to a relatively pure EV separation. EV membrane proteins CD63, CD81, and CD9 were then detected on the intact bdEV surface of brain regions by single-particle interferometric reflectance imaging sensor (SP-IRIS) (Fig. 3a) and multiplexed ELISA (Fig. 3b). All three surface proteins were detected above background on bdEVs from eight brain regions. For all regions, CD81 and CD9 were more abundant on the bdEV surface relative to CD63. By region, the level of CD63 was highest on bdEVs from CBLM followed by HIPPO. However, HIPPO and CBLM bdEVs also displayed more CD81 and CD9 than bdEVs from other brain regions.

**Figure 3.**
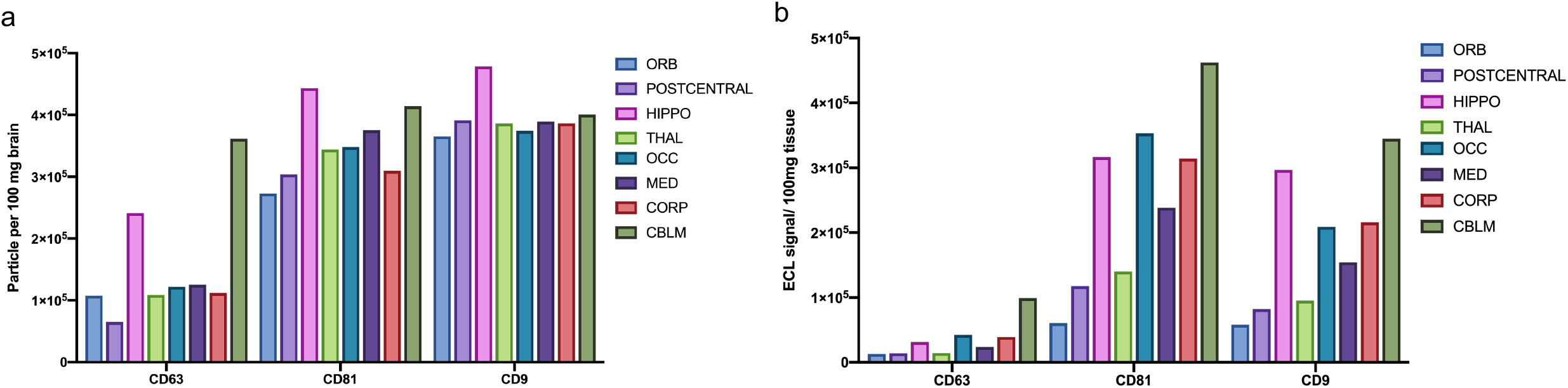
EV surface protein phenotyping. CD63, CD81, and CD9 were detected on the intact bdEV surface by single-particle interferometric reflectance imaging sensor (SP-IRIS) (a) and multiplexed ELISA (b) and normalized per 100 mg tissue input. bdEVs were captured by antibodies to EV membrane proteins and detected by signal from a cocktail of anti-tetraspanin antibodies (CD63, CD81, and CD9).

### Potential markers of cellular origin

To study the relative contribution of brain cell populations to bdEVs from different brain regions, 24 putative cell source markers (as listed in the Supplementary table 1) including 17 related to CNS cells (Fig. 4a) were assessed on the surface of intact bdEVs by multiplexed ELISA. The signal of each marker was normalized to the average signal of EV markers CD63, CD81, and CD9. Several differences were observed by marker and region (Fig.4b-f). Among putative neuron markers (Fig. 4b), NCAM, and CD90 were abundant on bdEVs from all regions, while CD166, CD24, CD271, and NRCAM were less abundant. The signal from some neuron markers was greater in bdEV populations from specific regions, e.g., NCAM signal in MED, CD271 in CBLM, and NRCAM in OCC. For microglia related markers (Fig. 4c), while HLA-DR/DP/DQ was least abundant on POSTC and HIPPO bdEVs, CD36 was in contrary most abundant on bdEVs from these two regions. In addition, the signal for astrocyte marker (Fig. 3d) CD44 was greatest on OCC and MED, and followed by THAL, CORP bdEVs compared with other regions. Among markers related to multiple CNS cell types (Fig. 3e), CD38 was greatest on OCC and CORP bdEVs, while CD15, TSPO, and GD2 were distributed evenly across regions. Several markers associated with immune cells and endothelia were also evaluated (Fig. 4f). Among them, the signal for endothelial marker CD29 and CD146 were greater, while CD307d and CD31 were almost undetectable. Prominently, CBLM bdEVs had the most CD29.

**Figure 4.**
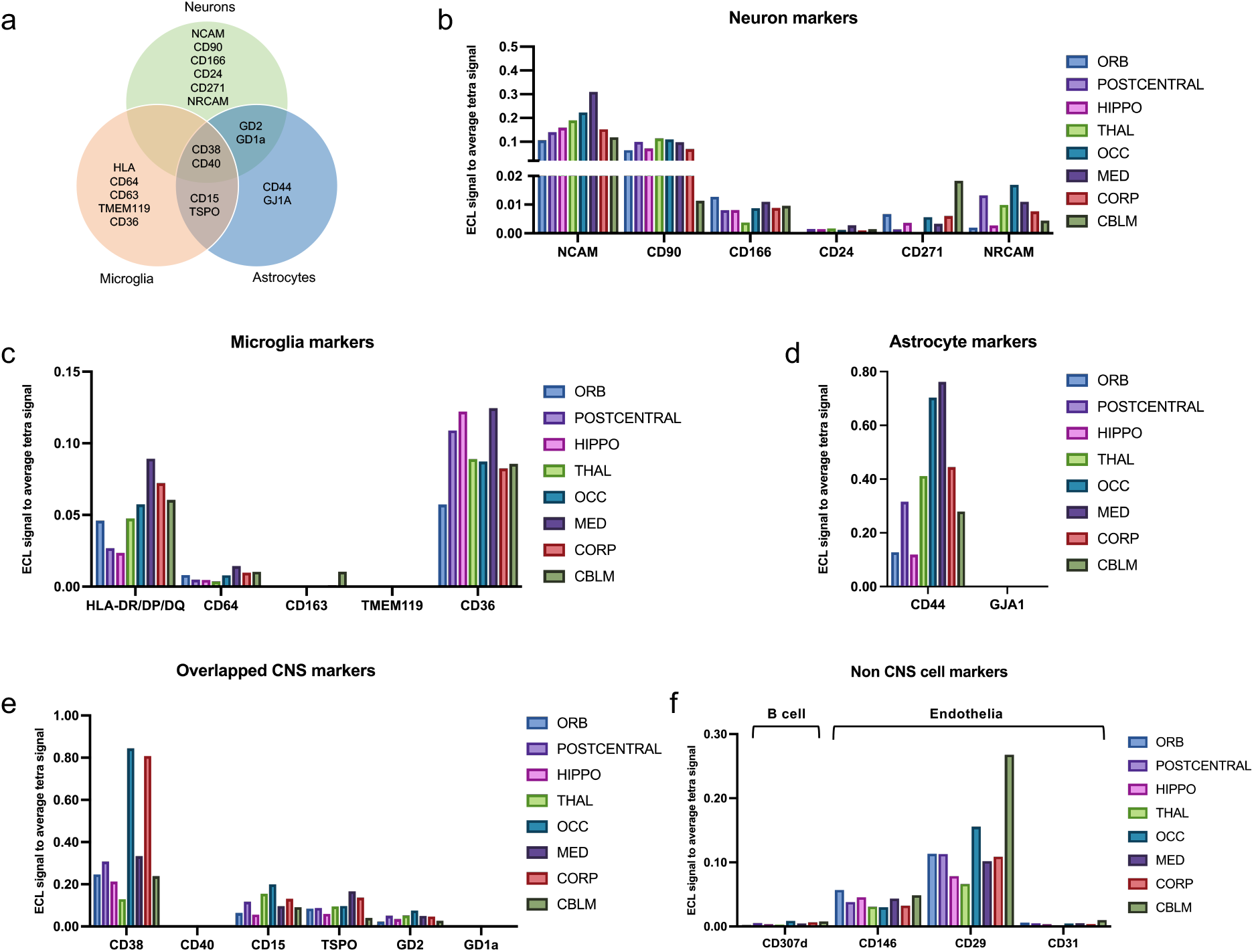
Cell-of-origin marker profile on the bdEV surface. (a) Distribution of markers by cell types: neurons, microglia, and astrocytes. Cell-enriched markers were used as bdEV capture antibodies; EVs were then detected by signal from a cocktail of anti-tetraspanin antibodies (CD63, CD81, and CD9). Levels of neuron (b), microglia (c), astrocyte (d), overlapping (e) and non-CNS cell (f) markers were then normalized to the average of tetraspanin capture spot signals.

### small RNA profiles

Small RNA (sRNA) sequencing of bdEVs from different brain regions yielded an average of 4.3M (± 3.7M) reads per sample (M = million, 1 × 10^6). After adapter clipping and removing reads shorter than 15 nt, 89.92% (± 2.4%) of bdEV reads mapped to the human genome (hg38). The percentages of reads mapped to various RNA biotypes are shown for bdEVs from different regions (Fig. S2). Reads mapping to rRNAs and messenger RNAs (mRNAs) were the most abundant sRNA biotypes in bdEVs (Fig. S2a), while reads mapping to vault RNAs, miRNAs, and pre-miRNAs were the least abundant (Fig. S2b). RNA biotype composition differences are shown for bdEVs from several regions (Fig. S2 a-c). For example, there were more mRNAs, miRNAs, pre-miRNAs, mtRNAs, and lncRNAs in CBLM, but more tRNAs and RNYs in MED.

Principal component analysis (PCA) of bdEV sRNA profiles clearly separated CBLM, THAL, and MED bdEVs from those of other brain regions (Fig. 5a). Focusing on miRNAs alone gave similar results. We identified the 20 miRNAs with the largest number of normalized counts in bdEVs from each region and then performed unsupervised clustering based on 15 miRNAs that were the most abundant across regions (Fig. 5b). Similar to the total Srna profile differences, CBLM bdEVs and THAL/MED bdEVs clustered apart from the others. Most of these miRNAs had greater counts in CBLM, OCC, THAL, and MED than in HIPPO, POSTC, CORP, and ORB (Fig. 5b). Furthermore, beyond these common miRNAs, bdEVs from certain regions were also enriched in specific miRNAs (Supplementary Table 2). For example, hsa-miR-137-3p and hsa-miR-744-5p were among the top 20 miRNAs in ORB but were not ranked within the top 20 in other regions.

**Figure 5.**
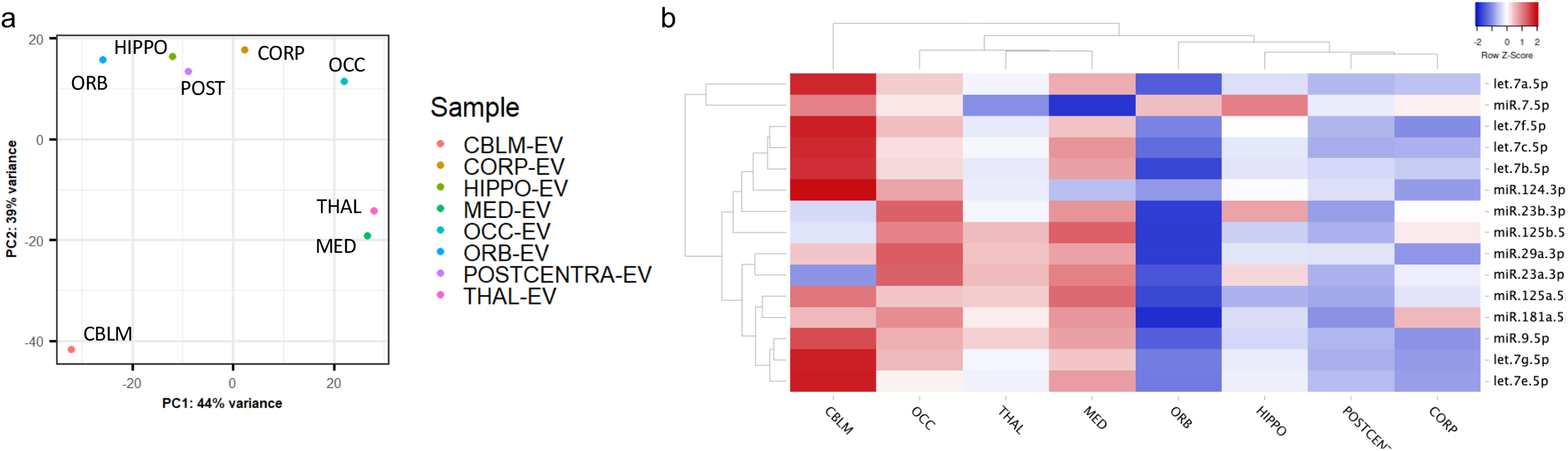
bdEV small RNA profiles. (a) Principal component analysis (PCA) based on quantitative small RNA profiles of bdEVs from different regions. (b) Unsupervised hierarchical clustering of 15 of the most abundant bdEV miRNAs across regions.

## Discussion

Studying bdEVs harvested from eight brain regions reveals evidence for region-specific differences in EV recovery, morphology, and molecular content. Although our findings are essentially a case study with regions from a single human brain, and thus do not support extensive discussion or speculation about the factors underlying apparent differences, our cell-enriched marker findings suggest that cell composition differences are likely to contribute. This study should now be expanded to assess whether or not the findings are reproducible, and, if so, if they hold across variables such as age, biological sex, diseases and disease stages, and species. Ultimately, building a regional “bdEV atlas” will assist with comparisons across studies that do not use the same anatomical region and spur new developments in region-specific monitoring and treatments.

## Figure legends

Figure S1 Western blots of ALIX, CD63, CD9, and calreticulin associated with BH and bdEVs. WB are representative of three independent human tissue EV separations from additional samples obtained from the brain bank for the purpose of protocol reproducibility assessment.

Figure S2 RNA biotypes of bdEVs from different regions. Percent of mapped reads for the RNA biotypes abundant across the regions (a), underrepresented across the regions (b), and other identified RNA biotypes in bdEVs (c).

## Acknowledgement

We thank members of the Witwer and Retrovirus Laboratories, Johns Hopkins University School of Medicine, and various members of the International Society for Extracellular Vesicles for valuable discussions and support. This work was supported in part by grants from the US National Institutes of Health: AI144997 (to KWW, with support for TAPD), MH118164 and AG057430 (to VM and KWW), UG3 and UH3 CA241694 (to KWW), UH2MH118167 (to DAR), supported by the NIH Common Fund, through the Office of Strategic Coordination/Office of the NIH Director. JHU Alzheimer ‘s Disease Research Centers NIH P30 AG 066507 and BIOCARD NIH U19AG033655.

