## Supplementary figures and images for "Towards a human brain EV atlas: Characteristics of EVs from different brain regions, including small RNA and protein profiles"

### Supplemental Figures

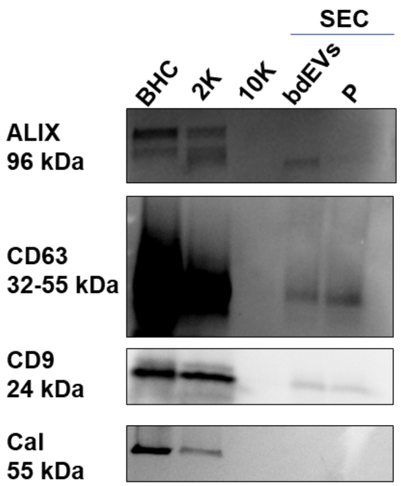

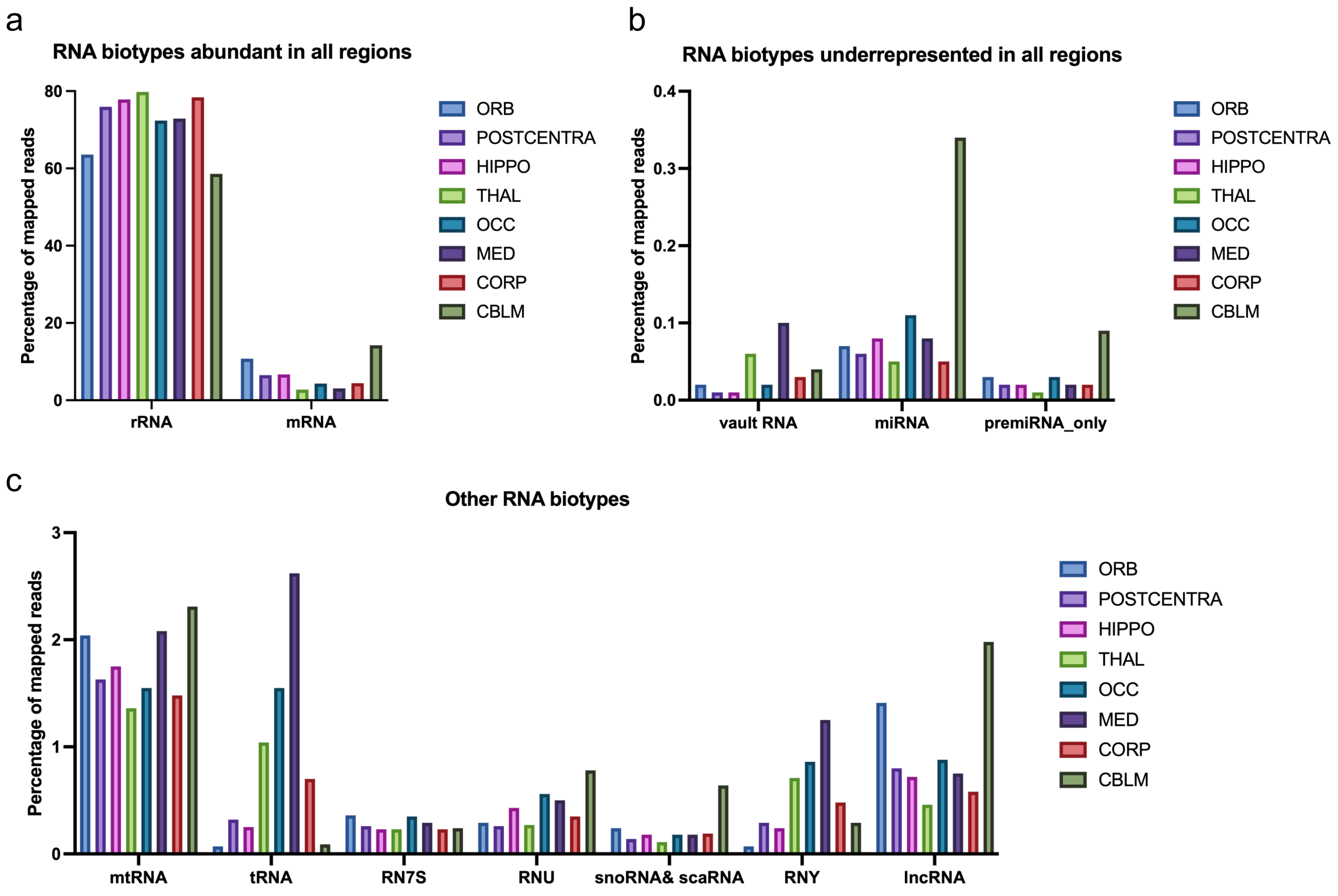
